# Structure-Based Design of Covalent Nanobody Binders for a Thermostable Green Fluorescence Protein

**DOI:** 10.1101/2024.11.22.624932

**Authors:** Zhihao Yue, Yanfang Li, Hongmin Cai, Hebang Yao, Dianfan Li, Aimin Ni, Tingting Li

## Abstract

Green fluorescence protein (GFP) lights up almost every aspects of life sciences. An ultra thermostable GFP (TGP), engineered from a coral GFP, offers potential advantages over traditional jellyfish-derived GFP due to its high stability. However, owing to its later discovery, TGP lacks the extensive toolsets available for GFP, such as heavy chain-only antibody binders known as nanobodies. In this study, we have structurally characterized TGP in complex with Sb92, a synthetic nanobody identified from a previous in vitro screening, revealing its precise three-dimensional epitope. This structural insight, alongside the previously characterized Sb44-TGP complex, have allowed us to rationally design disulfide bonds between antigen and antibody for tighter interactions. Using biochemical analyses, we identify two bridged complexes (TGP A18C-Sb44 V100C, TGP E118C-Sb92 S57C), with the TGP-Sb92 disulfide pair showing high resistance to reducing agents. Our study expands the toolkit available for TGP and should encourage its wider applications.

## Introduction

It is difficult to find a research field of biosciences without contribution of fluorescence proteins (FPs). In cell biology, FPs allow researchers to pinpoint protein locations (*1–3*), observe protein-protein interaction (*4–6*), and track biological processes at cellular (*7*), tissue (*8*), organ (*9*), and whole-body levels (*10*). In biochemistry, FPs enable convenient characterization of fusion partners including their expression and purification yield (*11*) and stability (*12–16*) without requiring pure samples. This is particularly beneficial for challenging-to-purify proteins, such as membrane proteins. FPs can also be engineered as reporters for cellular activities. For example, fusion proteins that combine circularly permutated green FP (GFP), calmodulin, and its binding peptide can track intracellular calcium changes by linking Ca²⁺-induced binding with improved GFP folding (*17*). Split FPs re widely used to detect protein-protein interaction (*18*), monitor inter-compartment protein translocation (*19*), and tether protein termini for crystallization (*20, 21*). Split GFP variants with inverted topology between the two halves have been used as sensors to detect specific protease activities, such as caspases, within cells (*22, 23*).

As a class, FPs are among the most extensively engineered proteins. The jellyfish GFP, for instance, has undergone modifications to improve brightness, maturation speed, folding efficiency, and color variation (*24*).

A particularly interesting GFP variant, Azami Green (*25*), originates from coral and has attracted considerable attention due to its unique properties. Evolutionarily distinct from jellyfish GFP, Azami Green shares only 27 % sequence identity with it and has a different chromophore, Gln-Tyr-Gly, compared to the Ser/Thr-Tyr-Gly in jellyfish GFP. Nevertheless, both retain a similar β-barrel structure known as a β-can (*26*). Due to its homology with red fluorescent proteins, Azami Green was the first GFP to be engineered to emit red fluorescence (*27*). By applying consensus mutagenesis, the Bradbury group created a variant named CGP (consensus green protein), which expressed with higher yield and brightness than Azami Green (*28*). Through recursive cycles of introducing and removing destabilizing elements combined with directed evolution, the same group developed an extremely stable CGP variant called eCGP123, which retains full fluorescence even after overnight heating at 80 °C (*29*). Additionally, eCGP123 maintains over 80% fluorescence in 6 M guanidine hydrochloride (GuHCl), a chaotropic condition typically disruptive to secondary structures. To address its tendency to aggregate, structure-guided mutations led to an even more stable variant, TGP (thermostable green protein) (*26*), which retains fluorescence after 50 days at 85 °C. TGP’s melting temperature (*T*_m_) is approximately 95 °C (with 20-min heating), about 20 °C higher than that of superfold GFP (*15*).

TGP has proven to be a superior fusion partner for membrane proteins, offering expression levels up to 30 times higher than conventional GFP fusions in widely used systems such as *Escherichia coli*, *Saccharomyces cerevisiae*, baculovirus, and mammalian cells (*15*). Its extreme stability also enables thermostability assessments of fusion membrane proteins across a wide temperature range using fluorescence-detection size-exclusion chromatography. In this approach, the fluorescence intensity of membrane protein-TGP fusions is plotted against temperature to determine the apparent *T*_m_. To facilitate affinity purification, our group previously (*15*) screened various synthetic nanobodies (known as sybodies) from mRNA libraries (*30*) using the highly efficient ribosome display method. While several structurally characterized nanobodies are available for the jellyfish GFP for use in protein purification (*31, 32*) and cell biology studies (*33–35*), such nanobodies for TGP has been lacking.

In this study, we have structurally characterized a sybody complexed with TGP, detailing the precise epitope and binding interactions. Using structural information, we engineered disulfide bonds to create covalent complexes between sybody and TGP. One resulting bridged complex showed substantial resistance to reducing agents. Our study expands the toolkit for TGP-based applications.

## Materials and Methods

### Molecular Cloning

Cysteine mutants were made by site-directed mutagenesis using standard polymerase chain reaction (PCR)-based protocols. Briefly, 20 ng of plasmids were used as PCR template, while mismatching primers were used to introduce intended mutations. PCR products were digested using *Dpn*I to remove template before purified using a clean-up kit (Cat. DC204, Vazyme). The resulted DNA were transformed into *Escherichia coli* DH5a cells. Single colonies with appropriate resistance from an agar plate were used to extract plasmids for Sanger sequencing.

### Purification of TGP and cysteine mutants

TGP and its cysteine mutants (A18C, E118C) were expressed in *E. coli* as a C-terminally His-tagged protein. BL21 (DE3) cells carrying the plasmid pETSG (***15***) was induced at an optical density at 600 nm (OD_600_) of 0.6–0.8 for 20 h at 20 °C using 1 mM isopropyl-β-_D_-thiogalactopyranoside (IPTG). Cells from 50 mL culture were suspended in a lysis buffer containing 150 mM NaCl, 50 mM HEPES pH 7.5, 1 mM PMSF and disrupted using an ultrasonic cell disruptor (Cat. JY92-IIN, SCIENTZ, CN) for 10-20 min in an ice-water bath. The lysate was centrifuged at 20,000 × g for 15 min at 4 °C. The supernatant was heated at 65 °C for 15 min, cooled in an ice-water bath, followed by centrifugation at 20,000 × g for 15 min at 4 °C to remove heat-induced precipitates. The supernatant containing heat-resistant TGP was incubated with 0.3 mL of Ni-NTA resin for 2 h with gentle agitation at 4 °C. The beads were loaded into a gravity column and washed with 20 column volume (CV) of lysis buffer supplemented with 30 mM imidazole. TGP was then eluted using 400 mM imidazole in the lysis buffer. TGP was quantified using absorbance at 280 nm measured on a Nanodrop machine with the theoretical molar extinction coefficient of 31,985 M^-1^ cm^-1^.

### Purification of sybody and cysteine mutants

Sybody and its Cys mutants were expressed in *E. coli* as a C-terminally His-tagged proteins (*15, 36*). MC1061 cells carrying the plasmid pSB_init_Sb44/Sb44 V100C or pSB_init_Sb92/Sb92 S57C was induced at OD_600_ of 0.6–0.8 for 20 h at 22 °C. Cells were lysed by abrupt osmotic shock as follows. Biomass from 1 L of culture was resuspended in 20 mL TES buffer (0.5 M sucrose, 0.5 mM EDTA, and 0.2 M Tris-HCl pH 8.0) for dehydration at 4 °C for 0.5 h. Dehydrated cells were rehydrated by diluting the cells with 40 mL of ice-cold MilliQ H_2_O at 4 °C for 1 h. Periplasmic extracts were collected by centrifugation at 20,000 × g at 4 °C for 30 min. The supernatant was adjusted to contain 150 mM of NaCl, 2 mM of MgCl_2_, and 20mM of imidazole. Two milliliter of Ni-NTA resin pre-equilibrated with 20mM imidazole in 150 mM NaCl and 20mM HEPES pH 7.5 were added into the supernatant for batch binding. The mixture was gently stirred at 4 °C for 1.5 h. The resin was packed into a gravity column and washed with 20 CV of 30 mM imidazole in 150 mM NaCl, 20mM HEPES pH 7.5. The sybody was eluted using 250 mM imidazole in the same buffer as above. Sybody proteins were quantified using absorbance at 280 nm measured on a Nanodrop machine. The molar extinction coefficient values of 32,457 M^-1^ cm^-1^ and 27,065 M^-1^ cm^-1^ were used for Sb44 and its mutant Sb44 V100C, and Sb92 and Sb92 S57C, respectively.

### Purification of sybody-TGP complexes

TGP (or its cysteine mutants) and sybody (or its mutants) were mixed at a molar ratio of 1:3 on ice for 20 min. The mixture was loaded onto a Superdex 200 Increase 10/300 GL column connected in a Bio-Rad NGC system equipped with a multi-wavelength absorbance detector in a running buffer containing 150 mM NaCl, 20 mM HEPES pH 7.5. Fractions were collected automatically by monitoring absorbance at 493 nm and 280 nm. Peak fractions were pooled for SDS-PAGE analysis.

### SDS-PAGE

Protein samples were mixed with equal volume of 2× Laemmli sample buffer with or without dithiothreitol (DTT). DTT concentrations were indicated in the maintext figures. Samples were loaded onto a home-made 12% gel for electrophoresis. Home-made GFP-tagged fluorescence markers (*37, 38*) were loaded alongside with broad-range standards. Gels were first imaged by in-gel fluorescence using a portable TGreen Transilluminator (Cat. No. OSE-470, Tiangen, Shanghai, China) with a smartphone. The same gel was then stained using Coomassie-blue for bright-field imaging.

### Crystallization

TGP for crystallization contained residues 1-218 of TGP with a SGGGSGGG linker between the C-terminal octa-histidine tag (*15*). TGP and Sb92 were mixed at a molar ratio of 1:1.2, and the complex was purified using size exclusion chromatography in a running buffer containing 150 mM NaCl, 20 mM HEPES pH 7.5. Pooled fractions were concentrated to 20 mg mL^-1^ using a 10-kDa cut off filtration membrane (Cat. UFC501096, Merck Millipore, Burlington, MA, USA). Sitting drop crystallization trials were set up by depositing 150 nL of the precipitant solution on top of 150 nL of protein with 70 μL of the reservoir using a Crystal Gryphon LCP robot (Art Robbins Instruments). Crystal plates were incubated at 20 °C in an incubator. Crystals grew in a precipitant solution containing 14.4 %(w/v) PEG 8000, 160 mM calcium acetate, 20% (v/v) glycerol, 80 mM sodium cacodylate / HCl pH 6.5.

### Data collection and structure determination

Crystals were harvested using a MiTeGen loop (Cat. M5-L18SP series, Ithaca, NY, USA) under an Olympus microscope (Model BX43F, Olympus, Tokyo, Japan). Cryo-cooling was achieved by rapid plunging crystals into liquid nitrogen. Crystals were screened by X-ray diffraction at beamlines 18U1 at the National Facility for Protein Science in Shanghai (NFPS) at Shanghai Synchrotron Radiation Facility. Diffraction data were collected with a 50 × 50 μm beam on a Pilatus 6M detector with oscillation of 0.5° and a wavelength of 0.97930 Å. Data were integrated using XDS (*39*), scaled and merged using Aimless (*40*). The structure was solved by molecular replacement using Phaser (*41*) with a TGP monomer and sybody of the Sb44-TGP structure (PDB ID 6LZ2) (*15*) as the searching models. The model was built with 2F_o_-F_c_ maps in Coot (*42*), and refined using Phenix (*43*). Structure was visualized in PyMol (*44*).

### Data availability

Atomic coordinates and structure factors for the reported TGP-Sb92 structure are deposited in the Protein Data Bank (PDB) under accession codes of 7CZ0.

## Results and Discussion

### Crystal structure of the Sb92-TGP complex

Disulfide bonds are the most frequent covalent tethers that strengthen protein-protein interactions. To develop strong binders for TGP, we aimed to create a disulfide-linked nanobody-TGP complex through structure-based design. Currently, there existed only one TGP structure in complex with a synthetic nanobody named Sb44 (*15*). To expand the templates available for rational design, we crystalized TGP in complex with a different synthetic nanobody, Sb92 (**Fig. 1a**), which was also obtained from our previous study (*15*), in a precipitant solution containing 14.4 %(w/v) PEG 8000, 160 mM calcium acetate, 20% (v/v) glycerol, 80 mM sodium cacodylate / HCl pH 6.5 at 20 °C.

**Figure 1.**
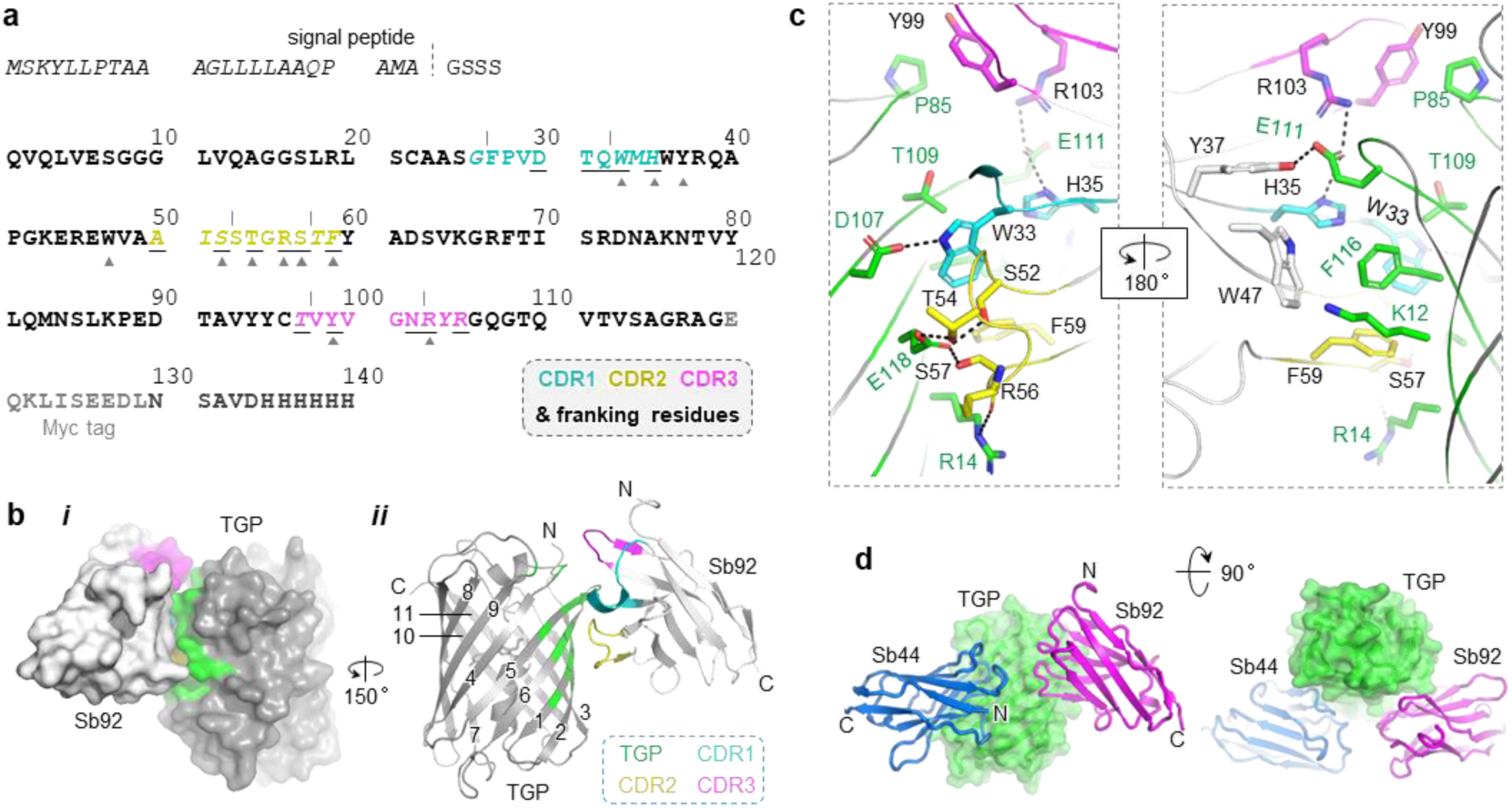
Structure of the Sb92-TGP complex. a Sequence of Sb92. CDR sequences (marked by vertical lines), along with flanking residues (italicized), are highlighted and color-coded as shown in the dashed box. Residues subject to randomization are underlined. Paratope residues are indicated by gray triangles beneath. Tag and signal peptide residues are shown in grey. A dash line marks the signal peptidase cleavage site. Signal peptide sequences are excluded from numbering. **b** Surface (**i**) and cartoon (**ii**) representation of Sb92 (light grey) bound with TGP (dark grey). TGP epitope residues are colored green. CDR1, CDR2, and CDR3 and their flanking residues are colored cyan, yellow, and magenta, respectively. Numbers indicate TGP β-strands. The N- and C-termini are labeled by N/C (**ii**). **c** Sb92-TGP interaction network. TGP residues are colored green. CDRs and their flanking residues of Sb92 are color-coded as in panel **a**. Framework residues are colored grey. Hydrogen bonds and salt bridges with distances within 3.2 Å are shown as black dashed lines. **d** Superposition of Sb44-TGP (PDB ID 6LZ2) and Sb92-TGP. CDR, complementarity-determining region; TGP, thermostable green fluorescence protein.

The crystals diffracted to 2.77-Å resolution with the *P*2_1_2_1_2 space group. The structure was solved by molecular replacement, using separate TGP and sybodies molecules (*15*) as search models, and refined to *R*_work_ and *R*_free_ values of 20.23 and 24.06%, respectively (**Table 1**). Each asymmetric unit contained four TGP-Sb92 complexes with high similarity, showing a Cα root mean square deviation (RMSD) ranging from 0.14 to 0.21 Å. Chain A (TGP) and chain E (Sb92) were used to describe the complex structure.

**Table 1.** Data collection and refinement statistics.

|  | Sb92-TGP |
| --- | --- |
| **Data collection** |  |
| Space group | *P*2<sub>1</sub>2<sub>1</sub>2 |
| Cell dimensions |  |
| *a*, *b*, *c* (Å) | 95.25, 130.15, 171.45 |
| *α*, *β*, *γ* (°) | 90, 90, 90 |
| Wavelength (Å) | 0.97930 |
| Resolution (Å) | 47.62 - 2.77 (1.85 - 2.77)<sup>a</sup> |
| *R*<sub>merge</sub> | 0.1384 (1.062) |
| *R*<sub>pim</sub> | 0.062 (0.466) |
| *I*/σ*I* | 9.2 (1.7) |
| Completeness (%) | 99.9 (99.9) |
| Multiplicity | 6.7 (7.0) |
| *CC* * <sup>b</sup> | 0.997 (0.936) |
| **Refinement** |  |
| Resolution (Å) | 47.62 - 2.77 |
| No. reflections | 54,791 |
| *R*<sub>work</sub> / *R*<sub>free</sub> | 0.2023/0.2406 |
| No. atoms | 10,620 |
| Protein | 10,440 |
| Ligand/ion | 180 |
| No. residues | 1,324 |
| B-factors (Å<sup>2</sup>) | 54.23 |
| Protein | 54.26 |
| Ligand/ion | 52.02 |
| R.m.s deviations |  |
| Bond lengths (Å) | 0.004 |
| Bond angles (°) | 0.715 |
| Ramachandran |  |
| Favoured (%) | 97.37 |
| Allowed (%) | 2.47 |
| Outlier (%) | 0.15 |
| **PDB ID** | 7CZ0 |

<sup>a</sup>Highest resolution shell is shown in parenthesis. <sup>b</sup> $CC^* = \sqrt{\frac{2CC_{1/2}}{1+CC_{1/2}}}$

### Molecular basis for Sb92-TGP binding

Sb92 binds to the TGP β-can at strands β1, β2, β5, and β6, with a buried surface area of 619 Å^2^ (**Fig. 1b**). All three complementarity-determining regions (CDRs), with CDR2 and CDR3 sandwiching CDR1, contributed to binding. In addition, the interactions involved two framework residues Tyr37 and Trp47 (**Fig. 1a**), a relatively common feature for nanobody-antigen interactions (*15, 45*).

The Sb92-TGP interaction involved hydrophobic packing, hydrogen bonding, and salt bridging. Specifically, the bulk side chain of Trp33 in Sb92 CDR1 inserts into a hydrophobic cage formed by Phe116’ and the hydrocarbon portion of Asp107’, Thr109’, and Glu118’ from TGP (TGP residues are indicated with a prime), as well as by Phe59 and Ser52 from Sb92 itself. The indole nitrogen of Trp33 forms a hydrogen bond with Asp107’. In CDR2, three hydroxyl-containing residues, namely Ser52, Thr54, and Ser57, participate in a hydrogen network with the acidic residue Glu118’ on TGP’s β6 strand, while Arg56 forms a hydrogen bond with the gualadine of Arg14’ via its backbone carbonyl group. In CDR3, Arg103 forms a salt bridge with Glu111’, which also interacts with H35 from CDR1 and Tyr37 from the framework region. Finally, Trp47 from the framework and Tyr99 from CDR3 pack against Phe116’ and Pro85’, respectively (**Fig. 1c**).

Sb92 was selected from a synthetic library designed to maintain a concave paratope shape (*30*). This library contains several features. Frist, the CDR regions, along with several flanking residues oriented away from the nanobody hydrophobic core, are randomized (**Fig. 1a**). Second, certain CDR residues are not randomized. Third, the library is designed to have a short CDR3 containing only 5 residues. The interaction profile remarkably aligns with these designed features. Specifically, randomized CDR-flanking residues (Trp33, His35, Ser52, Phe59, and Arg103) make a significant contribution for TGP-binding. Comparatively, the CDR regions include fewer residues overall, and none of the fixed residues in randomized regions contribute to the interaction. Finally, unlike natural nanobodies where CDR3 typically dominates antigen recognition, Sb92’s CDR3 plays a minor role in TGP interaction, involving only two residues (**Fig. 1a**, **Fig. 1c**).

The binding epitope of Sb92 on TGP differs from that of Sb44 (*15*), the only other structurally characterized sybody for TGP (**Fig. 1d**). When superimposed, Sb92 and Sb44 display a head-to-head configuration (**Fig. 1d**).

### Design and purification of cysteine mutants for intermolecular disulfides between sybody and TGP

To engineer disulfide bonds between sybodies and TGP, we analyzed the complex structure to identify residue pairs with Cβ distances within 5.0 Å. This effort identified TGP Ala18 with Sb44 Val100 (**Fig. 2a**), and TGP Glu118 with Sb92 Ser57 (**Fig. 2b**) as potential candidates for mutation to cysteine.

**Figure 2.**
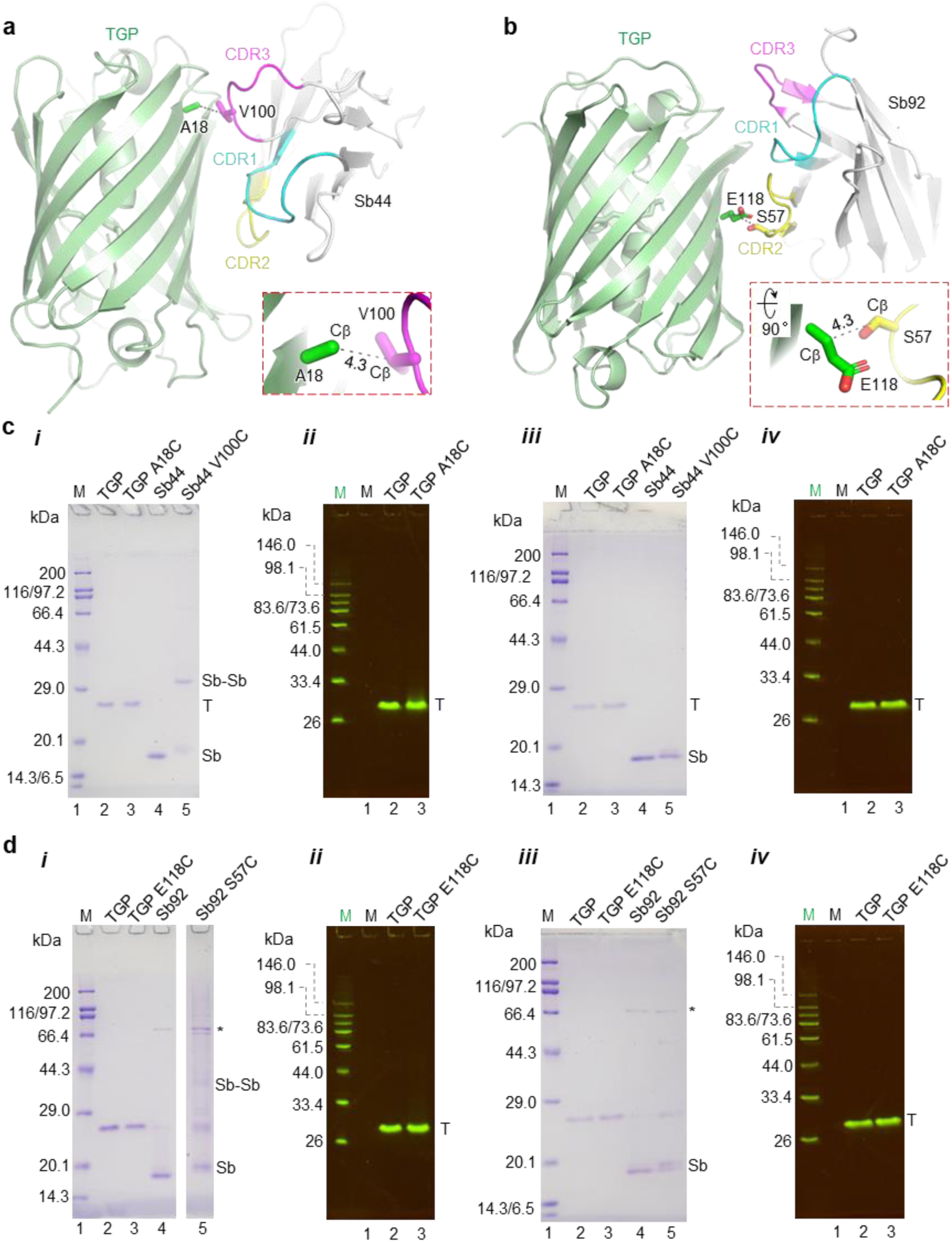
Structural Identification and Purification of Sybody-TGP Disulfide Mutation Candidates. **a, b** Cartoon representation of Sb44-TGP (**a**) and Sb92-TGP (**b**) complexes with candidate residues for cysteine mutations in stick representation. TGP and CDR regions are color-coded as shown. Cβ distances (Å) are indicated by numbers alongside black dash lines. **c, d** SDS-PAGE of TGP mutants with their respective Cys-pair mutants of Sb44 (**c**) and Sb92 (**d**). Gels were run under non-reducing (**i, ii**) or reducing conditions (**iii, iv**). Gels were visualized by Coomassie-blue staining (**i, iii**) or in-gel fluorescence (**ii, iv**). Home-made fluorescence standards (green M) and broad-spectrum markers (black M) were loaded on the left. Protein identities are labeled at the top of the gel, while lanes are numbered at the bottom. Lanes for the fluorescence marker (**i, iii**) and the non-fluorescence marker **(ii, iv**) were omitted. Bands corresponding to TGP (T), sybody (Sb), sybody dimer (Sb-Sb), and contaminant (*) are labeled on the right.

The cysteine pair mutants were generated by site-directed mutagenesis. When expressed individually in *E. coli*, TGP A18C and TGP E118C achieved yields of 32 mg L^-1^ and 36 mg L^-1^, respectively, comparable to the wild-type TGP yield of 30 mg L^-1^. Both mutants exhibited fluorescence brightness similar to the wild-type TGP, with relative fluorescence units (RFU) of 697 (A18C), 559 (E118C), and 657 (wild-type) at a concentration of 1 μg mL^-1^, as measured from a calibration experiment using our plate reader. These results indicate that the cysteine mutations did not disrupt TGP’s overall folding or fluorophore maturation. The sybody mutants also yielded quantities close to their respective wild-type forms (Sb44 V100C, 16 mg L⁻¹; Sb92 S57C, 2.5 mg L⁻¹, versus Sb44, 24 mg L⁻¹ and Sb92, 3.0 mg L⁻¹).

Since the cysteine mutations were introduced on the surface, we analyzed the homodimer-forming tendency of the individually purified mutants using SDS-PAGE under reducing and non-reducing conditions. In addition to Coomassie-blue staining, in-gel fluorescence was used to visualize TGP due to its stability and hence the ability to fluoresce on SDS-PAGE. Both TGP A18C and E118C migrated as monomers on SDS-PAGE under all conditions (**Fig. 2c**, **2d**).

In contrast to the wild-type Sb44, the Sb44 V100C mutant displayed a ∼32-kDa band under non-reducing conditions, indicative of dimer formation, which reverted to the monomeric 16-kDa band under reducing conditions (**Fig. 2c**, **2d**). Similarly, Sb92 S57C showed a higher molecular weight band that was more resistant to reducing agents than Sb44 V100C. These results suggest that while TGP mutants remain monomeric, sybody mutants tend to form disulfide-linked homodimers, likely due to the flexible positioning of cysteine residues in CDR loops compared to the β-strand-constrained cysteine residues in TGP. This flexibility may increase collision frequency, facilitating disulfide formation.

### Designed disulfides link TGP to sybodies

To examine whether heterodimeric complexes formed as designed, we mixed the cysteine mutants at a 1:3 molar ratio (TGP:sybody) and purified the resulting complexes via gel filtration. As a control, the wild-type TGP and sybodies, which were known to form complexes on gel filtration, were analyzed in parallel. The gel filtration profile was monitored by absorbance at both 280 nm for protein detection and 493 nm specific for TGP detection.

For the Sb44 V100C and TGP A18C mixture, a single A493 peak was detected at the same elution volume (V_e_) as the wild-type Sb44-TGP complex (**Fig. 3a**), indicating a stable sybody-TGP complex in solution. The normalized A280 and A493 traces for the main peak were superimposable (**Fig. 3a**), suggesting a homogeneous complex without noticeable excess of free TGP or sybody. In addition to the main A280 peak, two minor A280 peaks with a V_e_ of 22.4 mL and 23.7 mL were observed for both Sb44 V100C- TGP A18C and wild-type Sb44-TGP complexes (**Fig. 3a**), indicating Sb44 and the mutants formed homodimer in solution, possibly via hydrophobic interactions at the CDR regions for both Sb44 and Sb44 V100C, via disulfide linking for A18C.

**Figure 3.**
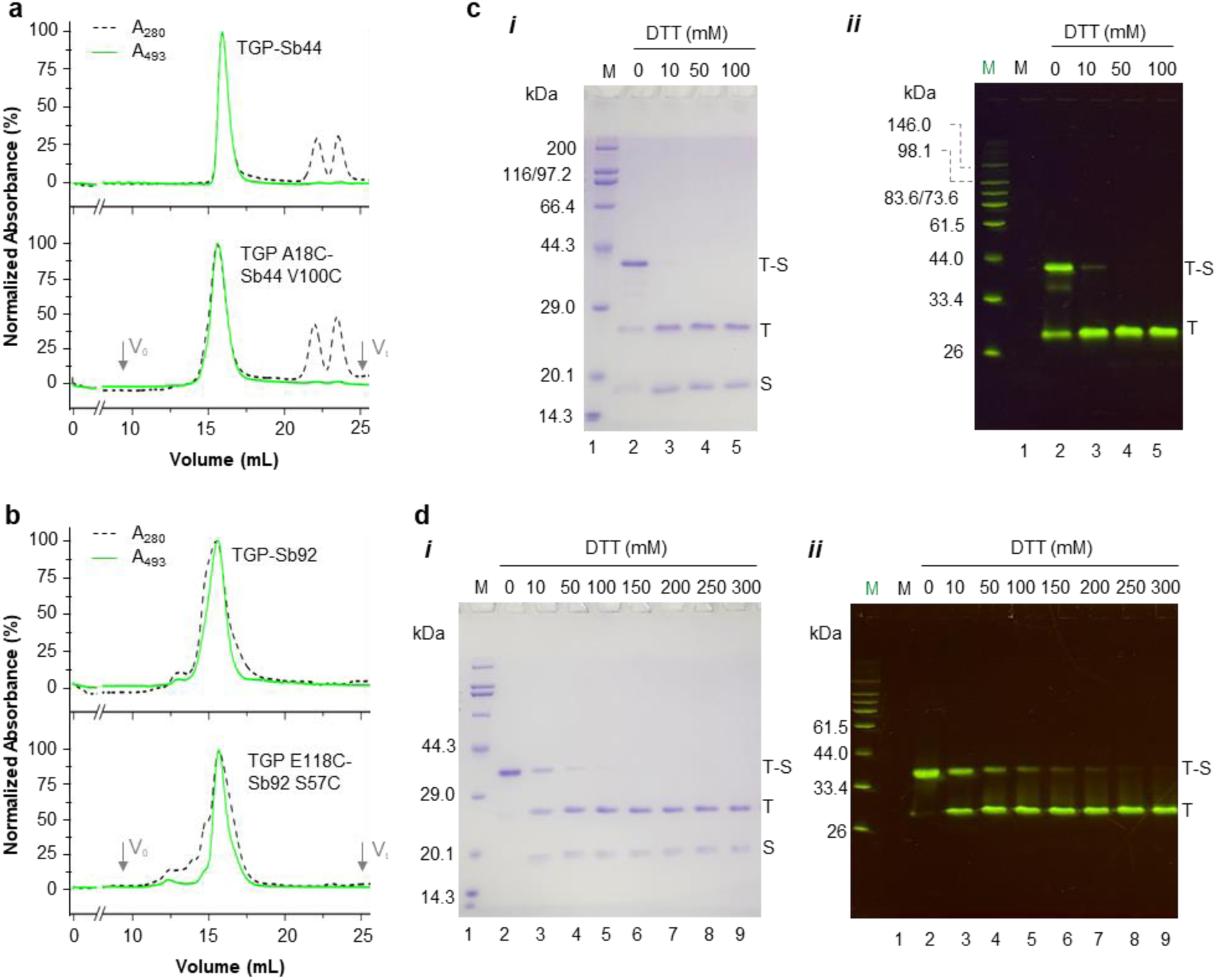
Purification and characterization of disulfide-linked sybody-TGP dimers. **a, b** Gel filtration of TGP A18C-Sb44 V100C (**a**) and TGP E118C-Sb92 S57C (**b**) alongside their respective wild-type complexes. Absorbance monitored at 493 nm (green, solid line) and 280 nm (black, dotted line) were normalized for plotting. Void and total volume are indicated by V_o_ and V_t_, respectively. **c, d** SDS-PAGE of the TGP A18C-Sb44 V100C (**c**) and TGP E118C-Sb92 S57C (**d**) complexes at varying dithiothreitol (DTT) concentrations indicated on top. Gels were visualized with Coomassie-blue staining (**i**) and in-gel fluorescence (**ii**). Home-made fluorescence standards (green M) and broad-range standards (black M) were included on the left. The lane for the fluorescence marker was not shown in **i**. Only relevant markers are labeled in **d**. Lane numbers are labeled at the bottom. T-S, T, and S indicate bands corresponding to TGP-sybody complex, free TGP, and free sybody, respectively.

For the mix of Sb92 S57C and TGP E118C, both the A280 and the A493 profile exhibited a major peak with minor shoulders to the left (**Fig. 3b**). The A493 peak was reasonably symmetric, suggesting the sybody-TGP complex was conformationally homogenous in solution. By comparison, the A280 peak was less symmetric, suggesting that Sb92 S57C forms some large homo oligomers that overlap with the heterodimer.

SDS-PAGE analysis under reducing and non-reducing conditions was performed on the peak fractions to confirm the presence of disulfide linkages in the sybody-TGP complexes. Under non-reducing conditions, the Sb44 V100C-TGP A18C complex appeared as a 40-kDa band on Coomassie-stained gels (**Fig. 3c**), corresponding to the disulfide-linked heterodimer, alongside two weaker bands corresponding to the free components. Addition of dithiothreitol (DTT) reduced the heterodimer band back to monomers, confirming the involvement of a disulfide bond. Notably, no homodimer band was observed for Sb44 V100C, indicating that its interaction with TGP outcompeted homodimer formation as seen in samples without the TGP cysteine pair (**Fig. 2c**).

For the Sb92 S57C and TGP E118C complex, a disulfide-linked heterodimer was also observed on SDS-PAGE. Unlike the Sb44 complex, no free monomers were detected, suggesting a more stable heterodimer. This complex was considerably more resistant to DTT; even with 0.1 M DTT in the loading buffer, residual heterodimer remained visible (**Fig. 3d**). Similar to Sb44 V100C, Sb92 S57C did not form detectable homodimers under these conditions (**Fig. 3d**).

Nanobodies and TGP are valuable tools in protein sciences and cell biology research, offering versatile applications in molecular recognition and imaging. In this work, we structurally characterized a synthetic nanobody, Sb92, in complex with TGP, revealing an atypical binding mode where CDR2, rather than the more commonly dominant CDR3 (*32, 46–50*), plays a significant role in antigen binding. Using this structural insight, together with data from the previously characterized Sb44-TGP crystal structure, we successfully designed disulfide-linked antibody-antigen complexes. To our knowledge, this represents the first example of a rationally designed covalent complex between a nanobody and its target.

The strong interactions achieved through the engineered disulfide pairs, particularly in the Sb92 S57C-TGP E118C complex, open new possibilities for trapping stable complexes under harsh conditions, including denaturing environments. One potential application is the purification and immunoprecipitation (IP) of TGP E118C-tagged proteins. Compared to conventional affinity purification, the covalent linkage approach developed here is expected to improve target protein purity by preventing complex dissociation during purification under stringent washing conditions. Moreover, the covalent complex could be used to capture interactions within the cells that are only transient or weak. For example, glycosylphosphatidylinositol-anchored proteins (GPI-APs) undergo several remodeling steps (*51*) in the endoplasmic reticulum membrane before export and secretion. Gaining insight into enzyme-substrate/product complexes along this pathway is crucial for mechanistic studies of GPI-AP quality control and sorting (*52*). One approach to stalling the remodelases with their GPI-AP products could involve tagging the enzyme and substrates with the Sb92 S57C-TGP E118C pair. The covalent linking would prevent the processed GPI-APs from entering the downstream export and secretion pipeline, thereby increasing the yield of intended enzyme-product complexes for structural studies. We are currently exploring these possibilities in our laboratory.

## Acknowledgements

We thank the staff scientists from BL18U1/BL19U1/BL10B beamline of National Facility for Protein Science in Shanghai (NFPS) at Shanghai Synchrotron Radiation Facility, for assistance during data collection. We thank Prof. Markus Seeger and Dr. Cedric AJ Hutter at University of Zurich, Switzerland for providing sybody libraries and technical guidance.

